# Shared characteristics establish *Corynebacterium glutamicum* ClpCP1P2 as a tractable model for mycobacterial orthologs

**DOI:** 10.64898/2026.11.12.766067

**Authors:** Pratistha Kandel, Farzaneh Hojjati, Karl R. Schmitz

## Abstract

Clp proteases are central to bacterial proteostasis, yet our biochemical understanding of actinobacterial Clp enzymes derives almost entirely from mycobacteria, where their essentiality complicates efforts to understand their physiological behavior. Here, we reconstitute and biochemically characterize the ClpCP1P2 protease from *Corynebacterium glutamicum*, a genetically tractable actinobacterium in which *clpC* and *clpX* are non-essential. We find that, as in *Mycobacterium tuberculosis*, ClpP1 and ClpP2 are individually inactive and form an active peptidase only when stabilized by ATP-bound ClpC or acyldepsipeptide antibiotics, with further stimulation by N-blocked peptide agonists. However, *C. glutamicum* ClpCP1P2 exhibits faster kinetics and greater conformational stability than its mycobacterial orthologs, proteolyzing full-length substrates rapidly without requiring peptide agonists. We additionally show that the anti-mycobacterial antibiotics rufomycin and cyclomarin A bind the *C. glutamicum* ClpC N-terminal domain, exert substrate-specific effects on proteolysis, and impair *C. glutamicum* growth. The ability of these compounds to elicit toxicity through a non-essential target suggests that they act via gain-of-function dysregulation rather than simple inhibition. Together, our results validate *C. glutamicum* as a tractable model for interrogating Actinomycetota Clp proteases.

**IMPORTANCE:** *Mycobacterium tuberculosis* remains one of the world’s deadliest bacterial pathogens and its Clp proteases are promising targets for new antibiotics. However, because these enzymes are essential in mycobacteria, it is difficult to study their physiological roles. Here, we show that the Clp protease of *Corynebacterium glutamicum*, a non-pathogenic relative in which the unfoldase ClpC is dispensable, shares the core assembly and regulatory features of its mycobacterial counterpart while being more stable and catalytically faster. Anti-mycobacterial ClpC-targeting antibiotics also act on this system, supporting a model in which they cause protease dysregulation rather than simple inhibition. These findings establish a genetically tractable model for dissecting actinobacterial Clp protease mechanisms and antibiotic action. They may also inform strategies for engineering regulated protein degradation in *C. glutamicum*, an important industrial microbe.

## INTRODUCTION

Virtually all bacteria possess Clp proteases that support general protein homeostasis and nuanced pathway regulation through the targeted degradation of cytosolic proteins. Clp proteases consist of a hexameric ring-forming AAA+ (ATPases associated with various cellular activities) unfoldase that recognizes protein substrates, unfolds them, and translocates them into a tetradecameric barrel-shaped peptidase for degradation (1,2). Decades of research on these enzymes – especially in *Escherichia coli* and *Bacillus subtilis* model systems – have informed detailed structural and mechanochemical models of their operating principles. Moreover, physiological studies have linked Clp proteases to virulence and/or pathogenesis in various disease-causing members of Pseudomonadota (synonym Proteobacteria; e.g. *Salmonella enterica* (3) and *Pseudomonas aeruginosa* (4)) and Bacillota (synonym Firmicutes; e.g., *Staphylococcus aureus* (5,6) and *Listeria monocytogenes* (7,8)). However, our understanding of Clp protease biochemistry outside of these phyla is limited.

Actinomycetota (synonym Actinobacteria), a diverse phylum including industrially, environmentally, and biomedically important species, possess notably complex Clp proteases. Most research on actinobacterial Clp proteases has focused on the globally important human pathogen *Mycobacterium tuberculosis*, where the strict essentiality of these enzymes makes them attractive antibacterial targets (9–11). While proteobacteria and firmicutes typically encode a single ClpP that forms a homomeric peptidase barrel, actinobacteria possess distinct ClpP1 and ClpP2 isoforms. In mycobacteria, biochemical and structural studies have established that these isoforms are individually inactive, but together assemble into a catalytically active ClpP1P2 heteromer composed of separate ClpP1 and ClpP2 rings (11,12). The ClpC1 or ClpX unfoldase docks coaxially atop the ClpP2 face of the peptidase barrel to form the full ClpC1P1P2 or ClpXP1P2 protease (13–15).

Mycobacterial Clp proteases also possess unusual assembly dynamics. Docking of Clp unfoldase to ClpP1P2 and binding of polypeptide to the peptidase catalytic sites stabilizes active ClpP1P2. Correspondingly, depletion of substrate (or of ATP, which is required for unfoldase binding) leads to disassembly of the active complex, which may serve as a layer of physiological regulation (16). Several compounds have been described that stimulate ClpP1P2 assembly and activity *in vitro*: N-blocked peptide agonists (e.g., carboxybenzyl-Leu-Leu, benzoyl-Leu-Leu) mimic substrate contacts by binding to peptidase active sites (12,17), while acyldepsipeptide antibiotics (ADEPs) mimic unfoldase contacts on the surface of the peptidase (15,17–20). It is not yet known whether these assembly characteristics are recapitulated by Clp proteases in other actinobacterial species.

While mycobacteria have served as *de facto* model systems for understanding actinobacterial Clp proteases, the essentiality of Clp enzymes in mycobacteria presents obvious experimental challenges. Depletion or inhibition of mycobacterial Clp enzymes leads to systemic pathway dysregulation and death (21), which makes it difficult to identify proteolytic substrates. Consequently, few *bona fide* physiological substrates have been reported to date (12,22–25), and the specific pathways responsible for the essentiality of mycobacterial Clp proteases remain elusive. Essentiality also constrains efforts to characterize Clp-targeting antibacterial compounds. For example, several cyclic peptide antibiotics (e.g., rufomycin and cyclomarin (26–29) potently kill mycobacteria by targeting the highly conserved N-terminal domain of ClpC1(30–32). However, investigations into their mechanism of action reveal complex proteomic perturbations that may mask the proximal proteolytic dysregulation induced by the compounds (33). An alternative actinobacterial model system in which Clp proteases function similarly but are non-essential would circumvent many of these limitations, but few studies have probed Clp proteases in actinobacterial species beyond mycobacteria. These enzymes have been examined in *Streptomyces coelicolor* (34), *Streptomyces lividans* (35), and the natural producer of ADEPs, *Streptomyces hawaiiensis* (36–38). However, *Streptomyces* express five ClpP isoforms whose levels are governed by complex regulation, and also possess an expansive proteolytic network that includes the proteases Lon, FtsH, and Mpa-20S (37,39).

By contrast, a simpler proteolytic landscape is found in *Corynebacterium glutamicum*, a genetically tractable microbe in the order Mycobacteriales that has been widely engineered for synthetic biology applications and industrial-scale biomanufacturing (40–43). Clp proteases are the only cytosolic AAA+ protease in *C. glutamicum*, which lacks orthologs of Lon and Mpa-20S, allowing more facile attribution of proteolytic programs to individual proteases. (The membrane-anchored protease FtsH is also present, but is thought to primarily process transmembrane and membrane-associated targets (44).) *C. glutamicum* Clp enzymes (ClpX, ClpC, ClpP1 and ClpP2) share 70 - 80% sequence identify with their *M. tuberculosis* orthologs, and studies have found that *clpC* and *clpX* are individually non-essential (45,46), enabling deletion and depletion studies that are impossible in mycobacteria. While prior studies have modulated *C. glutamicum* Clp protases in the context of protein expression and metabolic engineering (47–49), a detailed biochemical characterization of these enzymes has not been reported.

In this study, we reconstruct *C. glutamicum* ClpCP1P2 *in vitro* and examine its assembly, activity, and regulation. While we observe increased stability compared to mycobacterial counterparts, their major biochemical characteristics are similar, suggesting shared functional and regulatory paradigms across Actinomycetota. Interestingly, anti-mycobacterial ClpC-targeting antibiotics also dysregulate *C. glutamicum* ClpCP1P2 *in vitro* and impede cell growth. The fact that these compounds exert effects through a non-essential target aligns with recent findings (33) and suggests that they cause protease dysregulation rather than inhibition. Our work validates *C. glutamicum* as a tractable alternative to mycobacteria for investigating actinobacterial Clp proteases, and supports efforts to harness proteolytic pathways for nimble regulation of *C. glutamicum* pathways and improving biomanufacturing efficiency.

## RESULTS

### Interaction partners stabilize the active *C. glutamicum* ClpP1P2 peptidase *in vitro*

Like other actinobacteria, *C. glutamicum* encodes separate ClpP1 and ClpP2 paralogs within a bicistronic operon (46) (**Fig. 1A**). (Note that *clpP* ortholog numbering in *C. glutamicum* ATCC 13032 is reversed compared to *M. tuberculosis* H37Rv. To facilitate comparison to *M. tuberculosis* enzymes, we refer to *C. glutamicum* ClpP isoforms by the name of their closest *M. tuberculosis* ortholog: *C. glutamicum clpP1* as ClpP2 and *clpP2* as ClpP1.) In *M. tuberculosis*, the active form of the Clp peptidase is a heteromeric ClpP1P2 tetradecamer, which is stabilized by interactions with a AAA+ unfoldase partner (ClpC1 or ClpX) and translocated substrate. We therefore tested whether similar interactions are required to stimulate activity in *C. glutamicum* orthologs. In the absence of other interaction partners, ^Cgl^ClpP1 or ^Cgl^ClpP2, individually or in combination, were unable to hydrolyze a fluorogenic Pro-Arg-Met-aminomethylcoumarin (PRM-AMC) tripeptide (**Fig. 1B**). Inclusion of ^Cgl^ClpC without ATP had no stimulatory effect, nor did inclusion of ADP, but inclusion of ^Cgl^ClpC together with ATP stimulated PRM-AMC cleavage by ^Cgl^ClpP1P2 ∼10-fold (**Fig. 1B, 1C**), confirming that only the ATP-bound state of ^Cgl^ClpC productively activates ^Cgl^ClpP1P2. Similar patterns of stimulation were observed with a longer internally quenched decapeptide substrate (**Fig. S1A**). Prior studies with mycobacterial enzymes demonstrated that the peptidase activity of ^Mtb^ClpP1P2 is strongly stimulated by some N-blocked peptides that bind to peptidase active sites (12,16,50,51). Inclusion of the N-blocked peptide agonist benzoyl-Leu-Leu (Bz-LL) similarly increased ^Cgl^ClpCP1P2 peptidase activity ∼4-fold (**Fig. 1B**).

**Figure 1.**
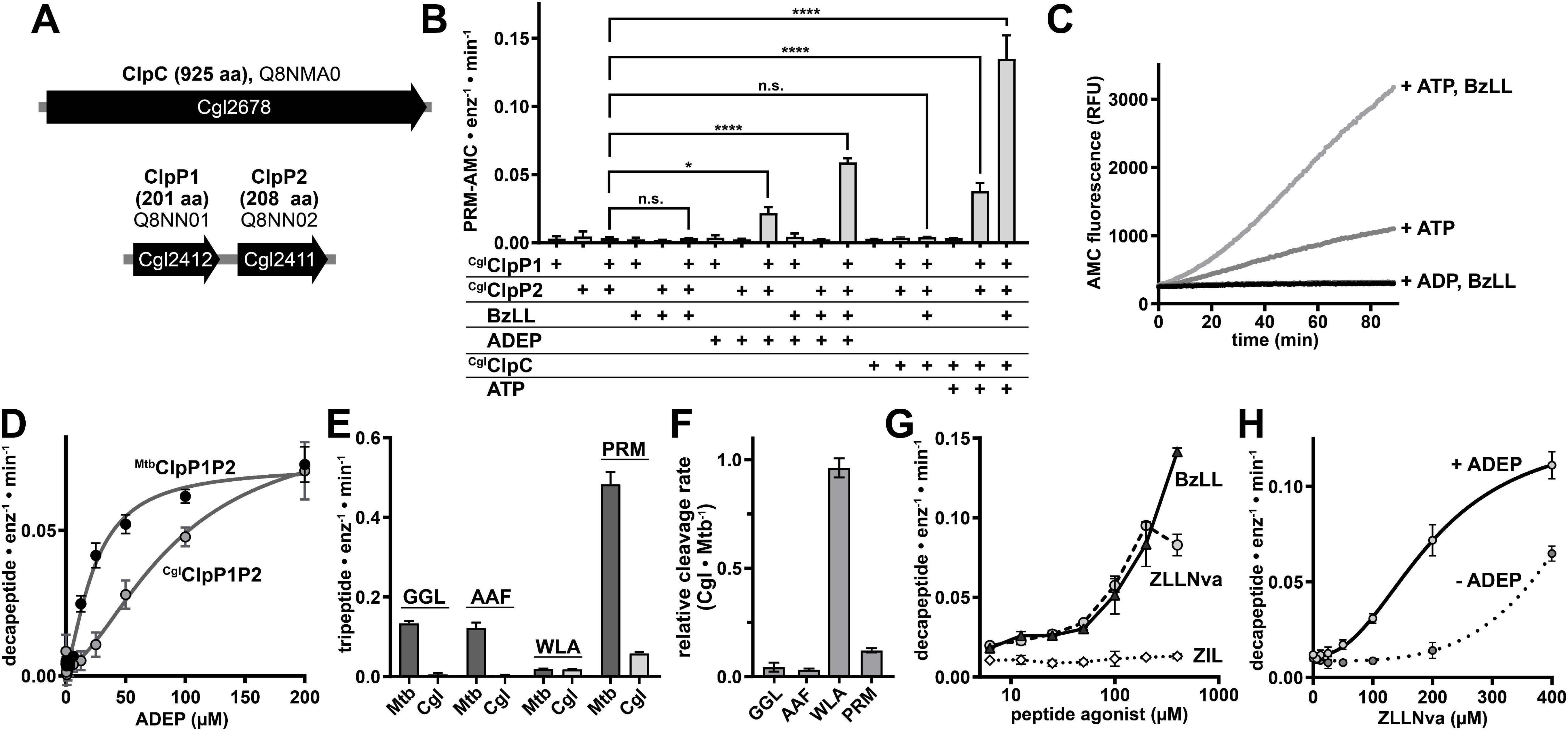
Unfoldase ClpC or ADEP and peptide agonist stabilize an active *C. glutamicum* ClpP1P2 peptidase *in vitro*. **A)** Operon organization of *C. glutamicum* loci encoding ClpP1, ClpP2 and ClpC. **B)** Hydrolysis of a fluorogenic Pro-Met-Arg-AMC peptide substrate (50 µM) was tested with combinations of ^Cgl^ClpP1 (0.25 µM) and/or ^Cgl^ClpP2 (0.25 µM) peptidase isoforms, agonist peptide Bz-Leu-Leu (100 µM), an ADEP antibiotic (100 µM), ^Cgl^ClpC (0.25 µM), and ATP (12.5 mM). Peptidase activity was only detected in the presence of ^Cgl^ClpP1 and ^Cgl^ClpP2, together with either ADEP or ^Cgl^ClpC with ATP. Inclusion of Bz-LL in either condition enhanced peptidase activity. **C)** Kinetic trajectories follow cleavage of 50 µM PRM-AMC by 0.25 µM ^Cgl^ClpP1P2 in the presence of 0.25 µM ^Cgl^ClpC with combinations of Bz-LL, ADP, or an ATP-regeneration system, illustrating that ATP is required for ^Cgl^ClpC-dependent stimulation of peptidase activity. **D)** ADEP dose-dependently stimulates cleavage of a decapeptide substrate (15 µM) by ^Mtb^ClpP1P2 (0.25 µM) or ^Cgl^ClpP1P2 (0.25 µM), with EC_50_ values of 24 ± 2 µM and 91 ± 20 µM, respectively. **E)** Assessment of peptidase activity by ^Mtb^ClpP1P2 (0.25 µM) or ^Cgl^ClpP1P2 (0.25 µM) towards a panel of tripeptide substrates, in the presence of Bz-LL and ADEP. **F)** Activities in E are shown as the ratio of ^Mtb^ClpP1P2 to ^Cgl^ClpP1P2 activity. **G)** Decapeptide (15 µM) cleavage by 0.25 µM ^Cgl^ClpP1P2 was measured across a range of concentrations of N-blocked peptides. **H)** Decapeptide cleavage by ^Cgl^ClpP1P2 (0.25 µM) was measured in the absence or presence of 100 µM ADEP across a range of concentrations of Z-Leu-Leu-Nva-CHO. Fit parameters are reported ± standard fit error. Error bars indicate SD among ≥3 replicates. Significance was determined by ordinary one-way ANOVA followed by Dunnett’s multiple comparison test, with the samples compared to CglClpP1P2 alone. Multiplicity-adjusted *p*-values are indicated as follows: * p ≤ 0.05, **** p ≤ 0.0001, n.s. non-significant.

Acyldepsipeptides (ADEPs), which structurally mimic the unfoldase Leu-Gly-Phe sequences that dock within hydrophobic pockets on the ClpP2 surface (15,17,20), can substitute for unfoldase to activate ^Mtb^ClpP1P2 (17). ^Cgl^ClpP1P2 was likewise activated by the combination of ADEP and agonist (**Fig. 1B, Fig. S1A**). Dose-response curves of ADEP stimulation (in the presence of Bz-LL) revealed relatively weak *EC_50_*values for both ^Cgl^ClpP1P2 (∼90 μM) and ^Mtb^ClpP1P2 (∼24 μM) (**Fig. 1D**), similar to values reported for ^Mtb^ClpP1P2 in prior studies (17,52). ADEPs, which are naturally produced by some *Streptomyces*, appear to have intrinsically weak affinity for actinobacterial ClpP1P2 enzymes.

We next compared the ability of ^Mtb^ClpP1P2 and ^Cgl^ClpP1P2 to cleave a panel of fluorogenic peptides (**Fig. 1E, 1F**). In line with its reported preference for Met, Leu or Phe in the P1 position (12,53), ^Mtb^ClpP1P2 demonstrated greatest activity towards PRM-AMC, moderate activity towards carboxybenzyl-Gly-Gly-Leu-AMC and Ala-Ala-Phe-AMC, and low activity towards Trp-Leu-Ala-AMC. ^Cgl^ClpCP1P2 was also most active toward PRM-AMC (although with ∼12% the activity observed with *M. tuberculosis* enzymes), weakly active towards AAF-AMC, and negligibly active against Z-GGL-AMC and WLA-AMC. The distinct preferences of ^Cgl^ClpCP1P2 suggest reduced ability to accommodate β-branched side chains.

We also examined dose-response patterns for agonist peptides. Structural studies of ^Mtb^ClpP1P2 show that agonists bind to peptidase active sites and stabilize the active state by preventing collapse into inactive conformations (17,50,54–56). Consequently, at sufficiently high concentrations, agonists are expected to competitively inhibit peptide cleavage, as has been reported with ^Mtb^ClpP1P2 (12,50). We tested the effects of the N-blocked agonists Bz-LL, Z-Leu-Leu-norvalinal (Z-LL-Nva-CHO) and carboxybenzyl-Ile-Leu (17,50) on ^Cgl^ClpP1P2 activity across a range of concentrations, in the presence of ADEP. While Z-IL produced no activation, Bz-LL and Z-LL-Nva-CHO both stimulated activity in a dose-dependent manner (**Fig. 1G**). Stimulation by Z-LL-Nva-CHO plateaued at 200 µM, likely due to limited solubility, similar to prior observations in mycobacteria (17). High concentration of Z-LL-Nva-CHO stimulated ^Cgl^ClpP1P2 peptidase activity even in the absence of ADEP, although with lower activity (**Fig. 1H**), echoing behavior reported with mycobacterial enzymes (50). Notably, mixtures of ClpP1 and ClpP2 from different orthologs were also enzymatically active in the presence of ADEP and agonist, suggesting that common interactions stabilize assembly (**Fig. S1B**).

### Agonist is dispensable for *C. glutamicum* ClpCP1P2 protease activity

Mycobacterial ClpC1P1P2 has been shown to proteolyze various model substrates *in vitro*, including FITC-casein (12), proteins carrying a C-terminal ssrA tag (Schmitz & Sauer, 2014), and fusion constructs incorporating the *M. smegmatis* 5-oxoprolinase pseudoenzyme MSMEI_3879 (23). We assayed proteolysis of ^CP7^GFP-ssrA, a circularly permuted GFP variant with reduced folded stability at the C-terminus (57,58), in reactions including combinations of ^Cgl^ClpC, ^Cgl^ClpP1, ^Cgl^ClpP2, and an ATP regeneration system (**Fig. 2A**). All three enzymatic components, along with ATP (**Fig. 2B**), were required for robust proteolysis, consistent with ATP-fueled proteolysis by the full ^Cgl^ClpCP1P2 complex. Degradation also occurred when we substituted ^Eco^ClpX for ^Cgl^ClpC (**Fig. S2A**), suggesting that common sequence elements (e.g., LGF/IGF-loops) stabilize engagement with ClpP1P2.

**Figure 2.**
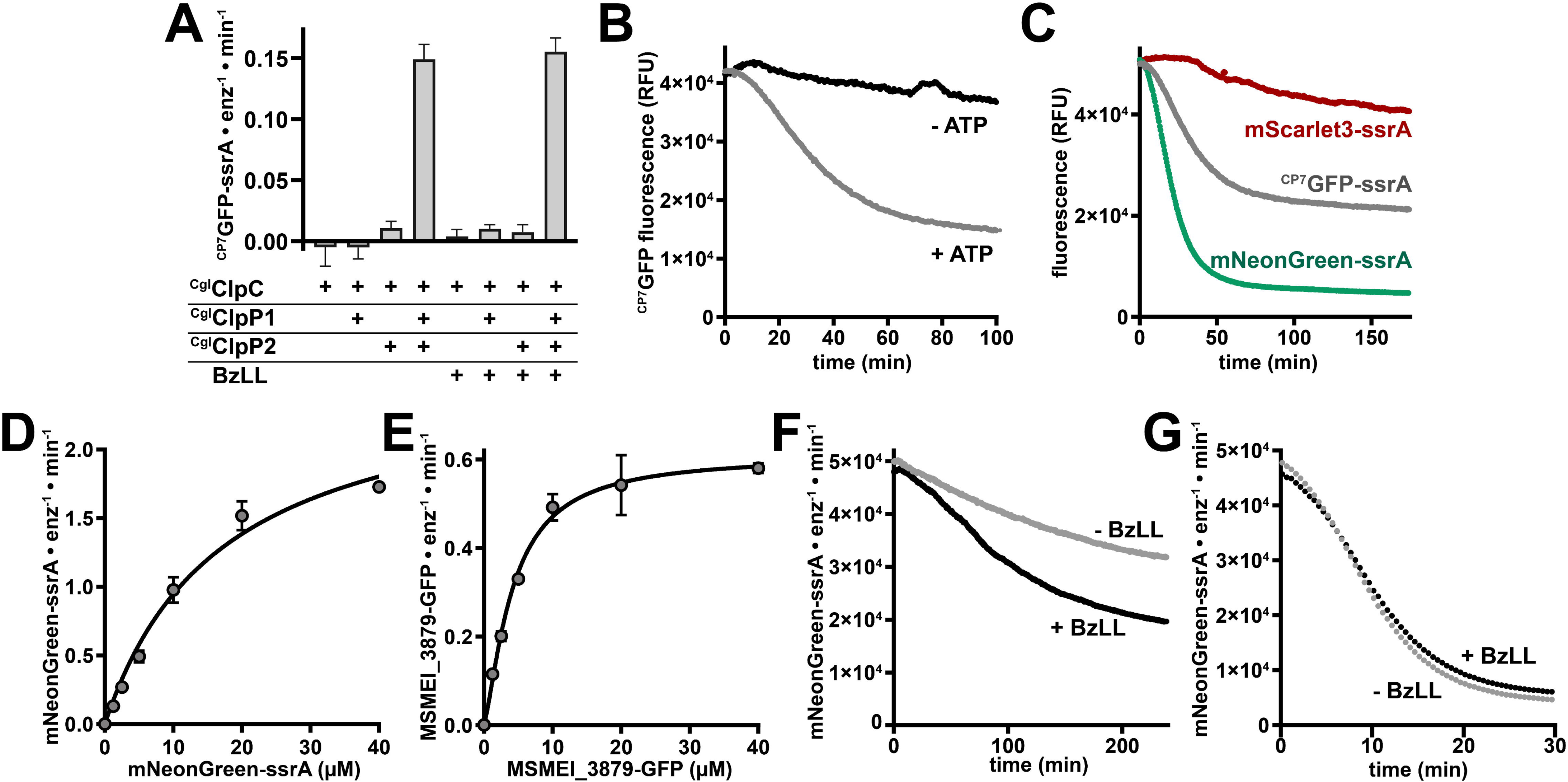
N-blocked peptides do not stimulate *C. glutamicum* ClpCP1P2 proteolysis. **A)** Degradation of a model ^CP7^GFP-ssrA substrate (20 µM) was tested with combinations of ^Cgl^ClpC (1 µM), ^Cgl^ClpP1 (0.5 µM), ^Cgl^ClpP2 (0.5 µM), and Bz-Leu-Leu (100 µM). Robust proteolysis required unfoldase along with both peptidase isoforms. **B)** Degradation of ^CP7^GFP-ssrA (20 µM) by 0.5 µM ^Cgl^ClpCP1P2 was monitored in the presence or absence of an ATP-regeneration system. **C)** Trajectories show degradation of the indicated model substrates (20 µM) by 0.5 µM ^Cgl^ClpCP1P2. **D)** Proteolysis of mNeonGreen-ssrA (20 µM) by ^Cgl^ClpCP1P2 (0.5 µM) was measured across varying substrate concentrations, and initial velocities were fit to the Michaelis-Menten equation (*K*_M_ = 17 ± 3 µM, *k*_cat_ = 2.6 ± 0.2 substrate•min^-1^•enzyme^-1^). **E)** Proteolysis of 20 µM MSMEI_3879-GFP by 0.5 µM ^Cgl^ClpCP1P2 was measured over varying substrate concentrations, and initial velocities were fit to the Michaelis-Menten equation (*K*_M_ = 5.1 ± 0.7 µM, *k*_cat_ = 0.68 ± 0.03 substrate•min^-1^•enzyme^-1^). Error bars reflect standard deviation among 3 technical replicates. Fit parameters are reported ± standard fit error. **F)** Proteolysis of 20 µM mNeonGreen-ssrA by 1.0 µM ^Mtb^ClpC1P1P2 was tested in the presence or absence of 100 µM Bz-LL. **G)** Proteolysis of 20 µM mNeonGreen-ssrA by 0.5 µM ^Cgl^ClpCP1P2 was assayed in the presence or absence of 100 µM Bz-LL.

While ^CP7^GFP-ssrA is a tractable substrate, it was only partially degraded over the reaction time course, mirroring incomplete degradation observed elsewhere for GFP-based substrates by ^Mtb^ClpC1P1P2 (24,59). We compared degradation kinetics of ^CP7^GFP-ssrA to ssrA-tagged variants of the fluorescent proteins mScarlet3 and mNeonGreen (**Fig. 2C**). While mScarlet3-ssrA was slowly and incompletely proteolyzed by ^Cgl^ClpCP1P2, mNeonGreen-ssrA was degraded ∼2.5-fold faster than ^CP7^GFP-ssrA and to a greater degree of completion. Assays with ^Mtb^ClpC1P1P2 revealed similar patterns of degradation (**Fig. S2B**), while *E. coli* ClpXP degraded mNeonGreen-ssrA about as well as ^CP7^GFP-ssrA (**Fig. S2C**). Michaelis-Menten analysis revealed that ^Cgl^ClpCP1P2 processes mNeonGreen-ssrA with >2.5-fold lower *K*_M_ and 30% greater *k*_cat_ than ^CP7^GFP-ssrA (**Fig. 2D, Fig. S2D, Table 1**), demonstrating that mNeonGreen-ssrA is a superior proteolytic substrate for *in vitro* assays with these enzymes. Additionally, we note that ^Cgl^ClpCP1P2 is a substantially faster protease than ^Mtb^ClpC1P1P2. ^Cgl^ClpCP1P2 degraded mNeonGreen-ssrA with 7-fold higher *k*_cat_ than ^Mtb^ClpC1P1P2, but similar *K*_M_ (**Fig. 2D**, **Table 1**). Similarly, ^Cgl^ClpCP1P2 proteolyzed MSMEI_3879-GFP with ∼2-fold higher *k_cat_* (**Fig. 2E**) than reported for ^Mtb^ClpC1P1P2 (23). The faster *in vitro* activities may reflect the faster growth rate of *C. glutamicum* compared to *M. tuberculosis*.

**Table 1:** Michaelis-Menten analysis of fluorescent substrate degradation by ^Mtb^ClpC1P1P2 and ^Cgl^ClpCP1P2.

|  | **<sup>Mtb</sup>ClpC1P1P2** |  | **<sup>Cgl</sup>ClpCP1P2** |  |
|---|---|---|---|---|
|  | *K*<sub>M</sub> (µM) | *k*<sub>cat</sub><br>(sub•min<sup>-1</sup>•enzyme<sup>-1</sup>) | *K*<sub>M</sub> (µM) | *k*<sub>cat</sub><br>(sub•min<sup>-1</sup>•enzyme<sup>-1</sup>) |
| **<sup>CP7</sup>GFP-ssrA** | - | - | 48 ± 7 | 2 ± 0.1 |
| **mNeonGreen-ssrA** | 11 ± 4 | 0.34 ± 0.04 | 17 ± 3 | 2.6 ± 0.2 |
| **MSMEI_3879-GFP** | 2.1 ± 0.4* | 0.26 ± 0.01* | 5.1 ± 0.7 | 0.68 ± 0.03 |

\*(23)

We also explored the dependence of proteolysis on agonist peptides. ZIL has been shown to enhance the activity of mycobacterial ClpC1P1P2 (16,50), suggesting that agonist helps stabilize the active proteolytic complex. Consistent with those results, we observed that Bz-LL stimulates mNeonGreen-ssrA degradation by ^Mtb^ClpC1P1P2 by ∼40% (**Fig. 2F and Fig. S2E**). By contrast, Bz-LL did not significantly alter proteolysis of mNeonGreen-ssrA by ^Cgl^ClpCP1P2 (**Fig. 2G and Fig. S2F**). The insensitivity of ^Cgl^ClpCP1P2 activity to agonist suggests that interactions made by the unfoldase partner and/or translocated substrate fully stabilize ^Cgl^ClpP1P2, demonstrating that the corynebacterial proteolytic complex is more stable than its mycobacterial counterpart.

### Rufomycin and cyclomarin A bind to ClpC and impair *C. glutamicum* growth

The non-ribosomally encoded peptide antibiotics rufomycin (Ruf) (27) and cyclomarin A (CymA) (26,28,29) bind to the N-terminal domain of mycobacterial ClpC1, dysregulate its activity, and cause mycobacterial cell death. To assess whether these compounds exert similar effects on *C. glutamicum*, we examined the effects of Ruf and CymA on growth across a range of concentrations. Both antibiotics caused impaired growth at concentrations ≥4 μM (**Fig. 3A and Fig. 3B**). To test whether antibiotics induce cell death, we treated *C. glutamicum* cultures with Ruf or CymA for varying durations and examined cell viability by spot plating. While most pre-incubation conditions had no influence on colony formation; 6 h pre-treatment with 8 µM of either antibiotic resulted in ∼10-fold reduction in apparent colony formation on agar plates after 24 h of growth (**Fig. S3A, S3B**). Notably, those samples eventually produced the same number of colonies as untreated controls after extending plate incubation to ∼48 h (**Fig. S3C, S3D**), suggesting that Ruf and CymA cause a growth delay in *C. glutamicum* rather than cell death.

**Figure 3.**
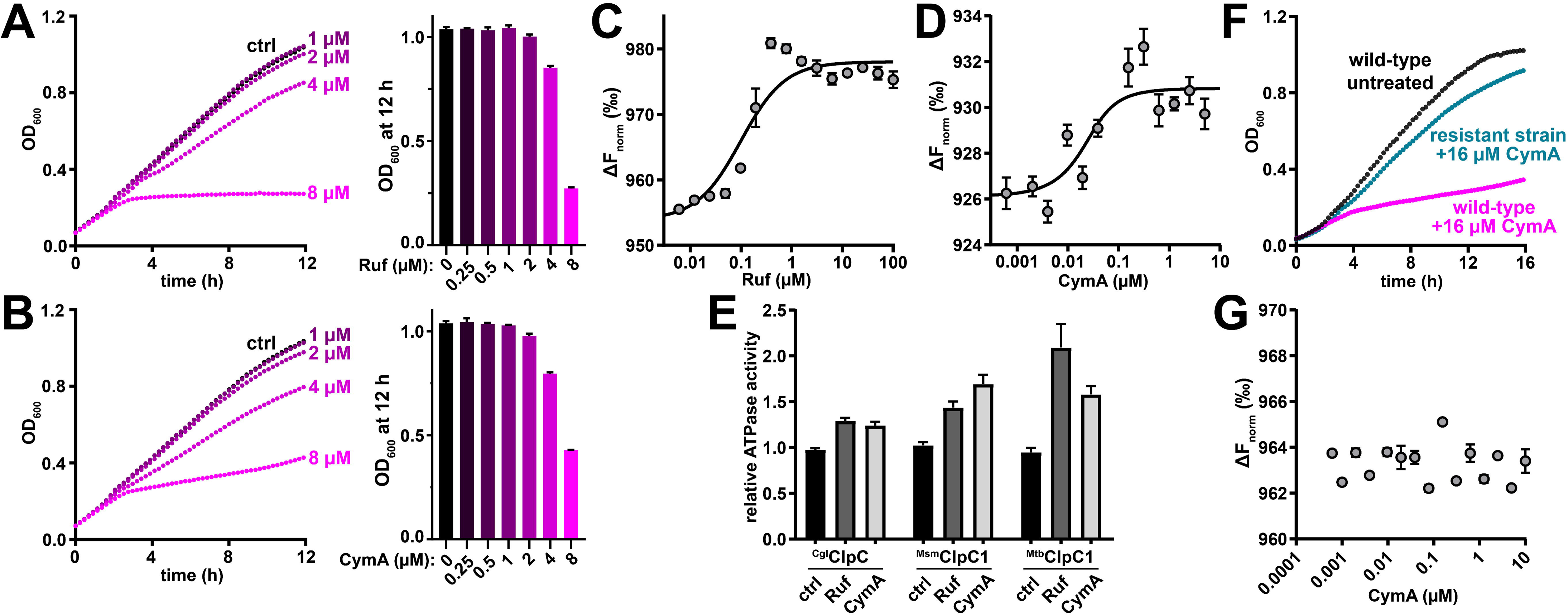
Rufomycin and cyclomarin A impede growth of *C. glutamicum* by targeting the N-terminal domain of ClpC. **A)** *C. glutamicum* cultures were treated with the indicated concentrations of Ruf, and growth was monitored by 600 nm absorbance. Representative growth curves are shown. Bar graphs show the average culture density after 12 h. **B)** *C. glutamicum* cultures were treated as in A, but with CymA. **C)** Binding of Ruf to purified ^Cgl^ClpC N-terminal domain was measured by microscale thermophoresis. Binding data was fit to a single-site binding model (*K*_D_ = 99 ± 23 nM) **D)** Binding of CymA to ^Cgl^ClpC N-terminal domain was measured as in C (*K*_D_ = 11 ± 8 nM). **E)** The ATPase rate of 1 µM ^Cgl^ClpC, ^Msm^ClpC1, or ^Mtb^ClpC1 was monitored in the absence or presence of 10 µM the indicated antibiotic. **F)** Representative traces show growth of untreated wild-type *C. glutamicum*, *C. glutamicum* treated with 16 µM CymA, or a resistant strain treated with 16 µM CymA. **G)** Microscale thermophoresis showed no binding of CymA to ^Cgl^ClpC N-terminal domain harboring an L88R mutation. Fit parameters are reported ± standard fit error. Error bars indicate standard deviation among ≥3 replicates.

We next sought to confirm that the toxicity of Ruf/CymA mechanistically arises through interactions with ClpC. The *C. glutamicum* ClpC N-terminal domain (NTD; residues 1-144) shares 88% sequence identity with the *M. tuberculosis* ClpC1 NTD, including residues critical for Ruf and CymA binding (eg, Met1, Phe2, His77, Phe80, and Glu89) (27,28,60). Microscale thermophoresis (MST) assays confirmed that Ruf and CymA bind to the ^Cgl^ClpC-NTD, with dissociation constants (*K*_D_ ≈ 90 nM and 10 nM, respectively) similar to the binding constants previously reported for *M. tuberculosis* (*K_D_* ≈ 63 nM and 4 nM with Ruf and CymA, respectively). (**Fig. 3C and Fig. 3D**) (60). Ruf and CymA also stimulated the *in vitro* ATPase rates of ^Cgl^ClpC, *M. smegmatis* ClpC1 and ^Mtb^ClpC1 to varying extents, consistent with prior reports (**Fig. 3E**) (26,61).

To confirm that ClpC is the specific molecular target, we exposed *C. glutamicum* cultures to increasing concentrations of CymA over a 7-day period and isolated surviving cells. These isolates grew normally in the presence of CymA (**Fig. 3F**). Whole-genome sequencing of two independent isolates revealed a common *clpC* polymorphism resulting in mutation of Leu88 to Arg. Strikingly, Leu88 lies within the CymA/Ruf binding site, and L88R mutations in *Mycobacterium abscessus*, *Mycobacterium marinum* and *Mycolicibacterium smegmatis* confer resistance to the structurally related antibiotic ilamycin (62). MST assays confirmed that CymA fails to bind to ^Cgl^ClpC-NTD^L88R^, confirming that CymA toxicity arises via ^Cgl^ClpC. Given that ClpC is non-essential in *C. glutamicum*, our findings align with recent studies in mycobacteria that implicate ClpC1 dysregulation, rather than outright inhibition, as the mechanistic cause of toxicity by ClpC1-targeting antibiotics (33).

### Rufomycin and cyclomarin A have substrate-specific effects on proteolysis

Studies with Ruf and CymA ^Mtb^ClpC1P1P2 have shown differing effects on proteolysis. Ruf is reported to inhibit proteolysis of FITC-casein and GFP-ssrA (23,27) while CymA stimulates degradation of these substrates (26). Recent studies have also revealed that CymA and ilamycin E (structurally similar to RufA) induce distinct proteomic perturbations in *M. tuberculosis* (33), suggesting nuanced substrate-specific dysregulation.

To begin to understand the species- and substrate-specific effects of Ruf and CymA on proteolysis, we tested the influence of each on the *in vitro* proteolytic activity of ^Cgl^ClpCP1P2 and ^Mtb^ClpC1P1P2 towards three model substrates: FITC-casein, mNeonGreen-ssrA and MSMEI_3879-GFP. Both antibiotics caused only minor inhibition of FITC-casein degradation by both enzymes (**Fig. 4A, 4B, Fig. S4A, S4B**), although Ruf shifted the proteolysis endpoint to a lower final fluorescence value, suggesting less complete degradation. Effects on mNeonGreen-ssrA varied by species. CymA stimulated increased proteolysis by ^Cgl^ClpCP1P2 (**Fig. S4C**), and both antibiotics caused ^Cgl^ClpCP1P2 to reach maximum degradation velocity earlier than a DMSO control (**Fig. 4C**), suggesting an accelerated conformational or assembly-dependent transition to the maximally active state. This effect was not observed with ^Mtb^ClpC1P1P2, for which antibiotics induced only mild inhibition (Ruf) or elevation (CymA) of the maximum degradation rate (**Fig. S4D**). Interestingly, both compounds induced strong and statistically significant inhibition of MSMEI_3879-GFP by both enzymes (**Fig. 4E, 4F, Fig. S4E, S4F**), with Ruf causing the greatest effect with ≥ 50% reduction in maximum degradation rate.

**Figure 4.**
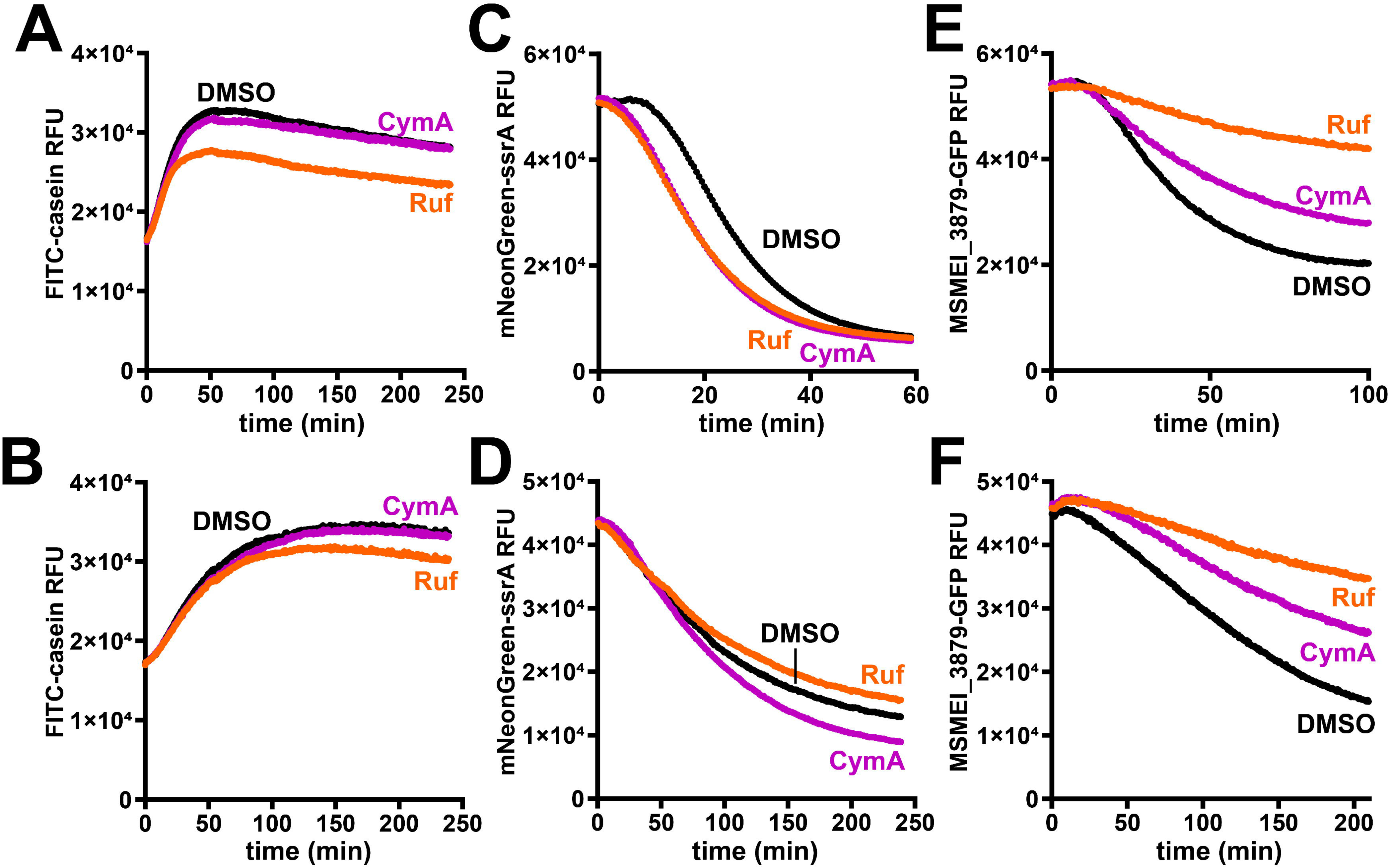
NTD-targeting antibiotics have substrate-specific effects on proteolysis. The influence of 10 µM Ruf, 10 µM CymA, or a DMSO control on proteolysis of FITC-casein (15 µM) by **A)** ^Cgl^ClpCP1P2 (0.5 µM) or **B)** ^Mtb^ClpC1P1P2 (1 µM); mNeonGreen-ssrA (20 µM) by **C)** ^Cgl^ClpCP1P2 (0.5 µM) or **D)** ^Mtb^ClpC1P1P2 (1 µM); or MSMEI_3879-GFP (10 µM) by **E)** ^Cgl^ClpCP1P2 (0.5 µM) or **F)** ^Mtb^ClpC1P1P2 (1 µM) was monitored by following changes in fluorescence. Traces are averaged from three biological replates. Corresponding proteolytic rates are shown in Fig. S4.

While the specific mode of recognition for these substrates is unknown, we previously speculated that MSMEI_3879 is recognized by interactions between the ClpC1 NTD hydrophobic groove (30,63,64) and an exposed hydrophobic surface on the MSMEI_3879 pseudoenzyme (23). Ruf and CymA bind directly to the NTD hydrophobic groove, and may therefore directly compete with MSMEI_3879 recognition. Overall, the fact that structurally similar antibiotics have differing effects on proteolysis across substrates and protease orthologs provides additional evidence to support the idea that NTD-targeting antibiotics have complex effects on protease activity state and substrate recognition, and are neither categorical inhibitors or stimulators.

## DISCUSSION

Although Clp proteases are widely distributed across Actinomycetota, our biochemical understanding of Clp protease orthologs across this diverse phylum comes almost exclusively from mycobacteria. Here, we report the *in vitro* reconstitution and biochemical characterization of the ClpCP1P2 protease from *C. glutamicum*, a non-mycobacterial Actinomycetota species. As in mycobacteria, ^Cgl^ClpP1 and ^Cgl^ClpP2 are individually inactive, with assembly of the functional heteromeric peptidase requiring stabilizing interactions to the LGF-binding pocket and/or peptidase active sites. This suggests that coupled assembly and activation are not a mycobacterial idiosyncrasy, and are likely a conserved regulatory paradigm across Mycobacteriales, and perhaps beyond. Moreover, our work establishes *C. glutamicum* as a tractable proxy for interrogating the mechanistic principles that govern mycobacterial Clp proteases specifically, and Actinomycetota orthologs more broadly.

Despite these overarching similarities, ^Cgl^ClpCP1P2 exhibits greater conformational stability and faster catalytic turnover than ^Mtb^ClpC1P1P2. In *M. tuberculosis*, peptidase and protease activities strongly depend on N-blocked peptide agonists, which mimic translocated substrate within the catalytic chamber, to shift the conformational equilibrium of ClpP1P2 toward its active state (12,16,17,50). By contrast, ATP-bound ^Cgl^ClpC or ADEP alone is sufficient to stimulate peptide cleavage by ^Cgl^ClpP1P2, and protein degradation proceeds readily without agonist supplementation. Thus, contacts from the docked unfoldase and translocating polypeptide suffice to stabilize active ^Cgl^ClpP1P2, reflecting a lower thermodynamic barrier to activation. Alongside the increased *k*_cat_ for protein degradation, this lower activation threshold may reflect evolutionary tuning to meet the demands of rapidly dividing *C. glutamicum* cells. Notably, we also found that mNeonGreen-ssrA is degraded faster and more completely than other fluorescent protein substrates by both corynebacterial and mycobacterial proteases, making it a superior model substrate for *in vitro* assays of actinobacterial Clp enzymes.

Our characterization of the ClpC NTD-targeting antibiotics Ruf and CymA provides physiological evidence that these antibiotics act through protease dysregulation rather than loss of function, in agreement with recent studies in mycobacteria (33). Both compounds bind the ^Cgl^ClpC NTD with nanomolar affinity and arrest *C. glutamicum* growth. Because *C. glutamicum clpC* is non-essential (45,46), simple inhibition of ClpC likely cannot account for growth impairment, indicating that antibiotic binding has toxic gain-of-function effects. Consistent with this, Ruf and CymA neither categorically inhibit nor uniformly stimulate ^Cgl^ClpCP1P2 and ^Mtb^ClpC1P1P2 *in vitro*. Rather, antibiotic effects likely reflect a combination of substrate-independent dysregulation (evidenced by perturbations in ATPase activity) and substrate-specific effects on recognition.

Together, these findings establish *C. glutamicum* as a biochemically faithful and genetically tractable model for investigating actinobacterial Clp proteases. The non-essentiality of *clpC* and *clpX* in *C. glutamicum*, as well as the absence of the AAA+ proteases Lon and Mpa-20S, facilitates Clp protease genetic knockouts, interactome trapping, and mechanism-of-action studies without the complexity or confounding lethality of mycobacterial models. An expanded understanding of these proteolytic machines will support development of Clp-targeting anti-mycobacterial drugs and enable synthetic biology approaches that harness Clp protease activity to dynamically tune metabolic flux in *C. glutamicum* biomanufacturing applications (47,49).

## MATERIALS AND METHODS

### Cloning of expression constructs

*M. tuberculosis* ClpC1 (23), mature ClpP1 (residues 7-200) (16), mature ClpP2 (residues 13-214) (16), *E. coli* ClpX (65), mNeonGreen-ssrA (55), ^CP7^GFP-ssrA (23), and MSMEI_3879-GFP (23) constructs were expressed and purified as described. DNA encoding constructs from *C. glutamicum* strain ATCC 13032, with *E. coli* codon optimization, was ordered from Twist Biosciences and cloned into pET vector backbones. Full-length ClpC (Cgl2678) was cloned in-frame with an N-terminal H_7_-SUMO tag. The ClpC N-terminal domain (residues 1-145) was cloned in-frame with a C-terminal sortase recognition site (LPETGG) followed by a H_6_ tag. Mature ^Cgl^ClpP1 (Cgl2412, annotated as ClpP2; Uniprot ID: Q8NN01; Met followed by residues 9-201) was cloned in-frame with a non-cleavable C-terminal H_6_ tag. Mature ^Cgl^ClpP2 (Cgl2411, annotated as ClpP1; Uniprot ID: Q8NN02; Met followed by residues 11-208) was cloned downstream of a self-cleaving RiboJ mRNA leader (66), to improve expression, and in-frame with a C-terminal Gly-Gly-Ser linker and non-cleavable H_6_ tag. ClpP1 and ClpP2 propeptide cleavage sites were inferred by comparison to *M. tuberculosis* orthologs. Codon-optimized DNA encoding mScarlet3 (67) was cloned in-frame with a C-terminal *M. tuberculosis* ssrA sequence (ADSNQRDYALAA) in a pET-derived plasmid. Constructs were verified by Sanger (Genewiz) or whole-plasmid sequencing (Plasmidsaurus or Genewiz).

### Protein expression and purification

Expression plasmids were transformed into a derivative of *Escherichia coli* strain ER2566 (NEB) harboring a *clpP* deletion. Cells were grown in 1.5 xYT (1:1 mixture of LB and 2xYT media supplemented with 1 mM MgSO_4_) at 37°C to exponential phase (OD_600_ ≈ 0.6-0.8). Expression was induced with 0.5 mM isopropyl-β-d-thiogalactopyranoside and cultures were incubated at 18°C overnight. Cells were harvested by centrifugation at 4000 × g for 30 min and resuspended in lysis buffer (25 mM HEPES, 10 mM imidazole, 500 mM NaCl, 10% glycerol, pH 7.5). 1 mM phenylmethylsulfonyl fluoride and 100 μL of Halt EDTA-free Protease Inhibitor Cocktail (Thermo) were added to ^Mtb^ClpC1, ^Eco^ClpX, ^Cgl^ClpC, ^Cgl^ClpC-NTD, and model substrate resuspensions. Cells were lysed by sonication and supernatant was collected after centrifugation at 10,000 × g for 45 min at 4°C. Recombinant proteins were purified from soluble lysate by Ni-NTA (MCLAB), anion-exchange (Source 15Q, Cytiva), and size-exclusion chromatography (Superdex 200, Cytiva). N-terminal H7-SUMO tags were removed prior to size exclusion chromatography by cleavage with *Saccharomyces cerevisiae* SUMO protease Ulp1. Stated concentrations of peptidase components reflect unfoldase hexamers or peptidase tetradecamers.

### Peptidase assays

Peptide cleavage assays included 0.25 μM ClpP1P2 in CPD buffer (25 mM HEPES, 200 mM KCl, 10 mM MgCl_2_, 0.1 mM EDTA, 1 mM DTT, pH 7.0). Experiments with fluorogenic tripeptides (Phe-Arg-Met-7-amino-4-methylcoumarin [AMC], Phe-Arg-Met-AMC, Z-Gly-Gly-Leu-AMC or Ala-Ala-Phe-AMC) included substrate at 50 µM, and cleavage was monitored by following fluorescence at 380 nm excitation and 460 nm emission. Experiments with internally quenched decapeptide (Abz-KASPVSLGY^NO2^D; Abz denotes the fluorophore 2-aminobenzoic acid and Y^NO2^ denotes the quencher 3-nitrotyrosine) included the substrate at 15 µM, and cleavage was monitored by following increase in 420 nm fluorescence upon 320 nm excitation. N-blocked agonist peptides (benzoyl-Leu-Leu, carboxybenzyl-Leu-Leu-norvalinal, or carboxybenzyl-Ile-Leu) and acyldepsipeptide (ADEP) were included at 100 µM, unless otherwise stated. ^Cgl^ClpC was included at 0.25 µM, along with 12.5 mM ATP and an ATP regeneration system consisting of 16 mM creatine phosphate and 0.32 mg/mL (w/v) creatine phosphokinase; assays incorporating ^Cgl^ClpC were pre-incubated at 30°C for 30 min before addition of peptide substrate. DMSO was included at a final concentration of 5% in all samples. Assays were carried out in 40 µL reactions in a black non-binding 384-well plate (Corning) at 30°C in a Tecan Spark plate reader.

### Proteolysis assays

*In vitro* proteolysis assays included unfoldase (1 μM ^Mtb^ClpC1, 0.5 μM ^Cgl^ClpC, or 1 μM ^Eco^ClpX), peptidase (0.5 μM ^Mtb^ClpP1P2 or 0.25 μM ^Cgl^ClpP1P2), substrate (20 µM ^CP7^GFP-ssrA, 20 µM mNeongreen-ssrA, 20 µM mScarlet-ssrA, 10 µM MSMEI_3879-GFP, or 15 µM FITC-casein [Sigma]), and an ATP regeneration system (12.5 mM ATP, 16 mM creatine phosphate, and 0.32 mg/mL (w/v) creatine phosphokinase) in CPD buffer. Assays were performed in 40 µL reactions in black 384-well plates at 30°C in a Tecan Spark plate reader. 100 μM Bz-LL was included where indicated. Ruf and CymA were added to 10 μM final concentrations. All reactions contained 5% (v/v) final DMSO. Degradation of fluorescent proteins followed by reduction in fluorescence over time at the following excitation/emission wavelengths: 380 nm/511 nm for ^CP7^GFP-ssrA and MSMEI_3879, 485 nm/530 nm for mNeonGreen-ssrA, and 570/595 nm for mScarlet3-ssrA. FITC-casein degradation was quantified by tracking the increase in fluorescence over time at 485 nm/538 nm.

### ATP hydrolysis assay

ATP hydrolysis assays was performed using 1 µM_6_ ^Cgl^ClpC or ^Mtb^ClpC1 and an NADH-coupled regeneration system (187.5 U/mL pyruvate kinase, 214.5 U/mL lactate dehydrogenase, 10 mM NADH, 50 mM phosphoenolpyruvate, 10 mM ATP) in CPD. Assays were carried out in 40 µL reactions at 30°C in clear non-binding 384 well plates (Corning) in a Tecan Spark plate reader. The oxidation of NADH to NAD^+^ was monitored via increase in 340 nm absorbance.

### Microscale Thermophoresis binding assay

^Cgl^ClpC-NTD bearing a C-terminal H_6_ tag was diluted to 100 nM in CPD buffer containing 0.05% (v/v) Tween20 and 5% (v/v) DMSO and incubated with HIS Lite™ OG488-Tris NTA-Ni Complex dye (AAT Bioquest) for 30 min at room temperature in the dark. 2-fold serial dilutions of Ruf and CymA were prepared in the same buffer conditions, along with a ligand-free control sample; mixed 1:1 with labeled ^Cgl^ClpC-NTD; and loaded into capillary tubes (World Precision Instruments). Thermophoresis measurements were collected on a NanoTemper Monolith NT.115 instrument at 40% LED and laser power at 25°C. Data were fit to single-site binding equation in GraphPad Prism.

### C. glutamicum growth curves

Wild-type *C. glutamicum* (carrying an empty Kan-resistant pZ8 plasmid(68)) was grown in 1.5 xYT medium at 30°C with constant 250 rpm orbital shaking to exponential phase (OD_600_ ≈ 0.6). The starter culture was used to inoculate 150 μL samples in fresh media in a 96-well plate at a starting OD_600_ of 0.05. A range of concentrations of Ruf or CymA was included; cells without added antibiotics served as negative controls. The outer rim of the plate was filled with 200 μL of sterile water and the plate was covered with a clear lid to maintain humidity. Absorbance was monitored over a duration of 12 h at 30°C in a Tecan Spark plate reader.

### Spot-plating viability assay

Wild-type *C. glutamicum* (carrying empty pZ8 plasmid)(68) starter cultures were prepared in 1.5 xYT medium, grown at 30°C to mid-log, and sub-cultured into 5 mL samples in 15 mL culture tubes at a starting OD_600_ of 0.05. 2-fold dilution series of CymA and Ruf was added to the samples, and samples were incubated at 30°C for 6 h. At 2 h intervals during incubation, samples were harvested, spun down at 4000 g for 5 min, and resuspended into fresh, antibiotic-free media. The recovered cells were serially diluted into sterile 1.5 mL tubes, plated in equal volumes onto LB agar, and incubated at 30°C for 48 h.

### Inducing acquired mutation of *C. glutamicum* cells against CymA

Wild-type *C. glutamicum* (carrying empty pZ8 plasmid(68)) was grown in 1.5 xYT to exponential phase and normalized to an OD_600_ of 0.05. 4 µM CymA was added, and the culture was allowed to grow for ∼18 h at 30°C. The culture was passaged daily at a 1:100 dilution into fresh media containing sequentially increasing doses of CymA (6, 8, 10, 12, 14 and 16 µM) over 6 days. At the end of 7^th^ day, the adapted cells were harvested by centrifugation at 4000 g for 5 min, washed to remove any residual antibiotic, plated into CymA-free LB agar, and incubated for 48 h at 30°C. Isolated colonies, and wild-type control colonies, were picked and cultured in liquid media. Genomic DNA was extracted using ZymoBiomics DNA Miniprep Kit and analyzed by whole genome nanopore sequencing (Plasmidsaurus).

## ACKNOWLEDGEMENTS

We thank members of the Schmitz lab for helpful advice and discussions, and J. Buehler for help with protein purification. We also thank S. Cho and S. Franzblau of the Institute for Tuberculosis Research at the University of Illinois Chicago for their gift of rufomycin, and F. Fei and J. Sello at UCSF for their gift of cyclomarin A. K.R.S. was supported by award R01AI171196 from NIH NIAID and NIGMS.

