## Supplemental Information - Fig. S1 - S4 for "Shared characteristics establish *Corynebacterium glutamicum* ClpCP1P2 as a tractable model for mycobacterial orthologs"

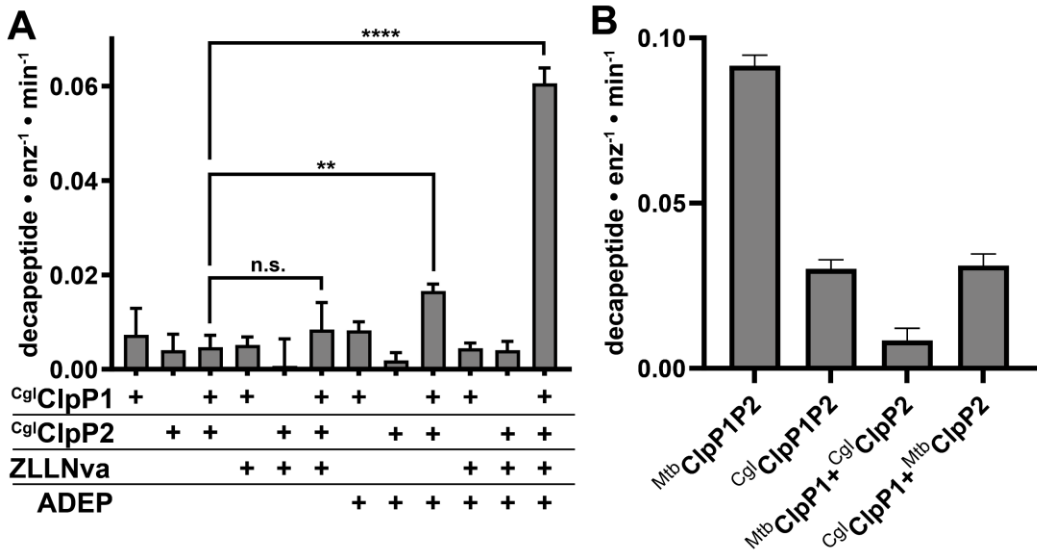

**Figure S1. Peptidase activity of *C. glutamicum* ClpP1P2. A)** Cleavage of a fluorogenic decapeptide substrate (15 μM) was tested with combinations of $^{Cgl}$ClpP1 (0.25 μM), $^{Cgl}$ClpP2 (0.25 μM), Z-Leu-Leu-Nva-CHO (100 μM), and ADEP (100 μM). **B)** Decapeptide (15 uM) cleavage was tested with combinations of $^{Mtb}$ClpP1 (0.25 μM), $^{Mtb}$ClpP2 (0.25 μM), $^{Cgl}$ClpP1 (0.25 μM), and $^{Cgl}$ClpP2 (0.25 μM) in the presence of Z-Leu-Leu-Nva-CHO (100 μM) and ADEP (100 μM).

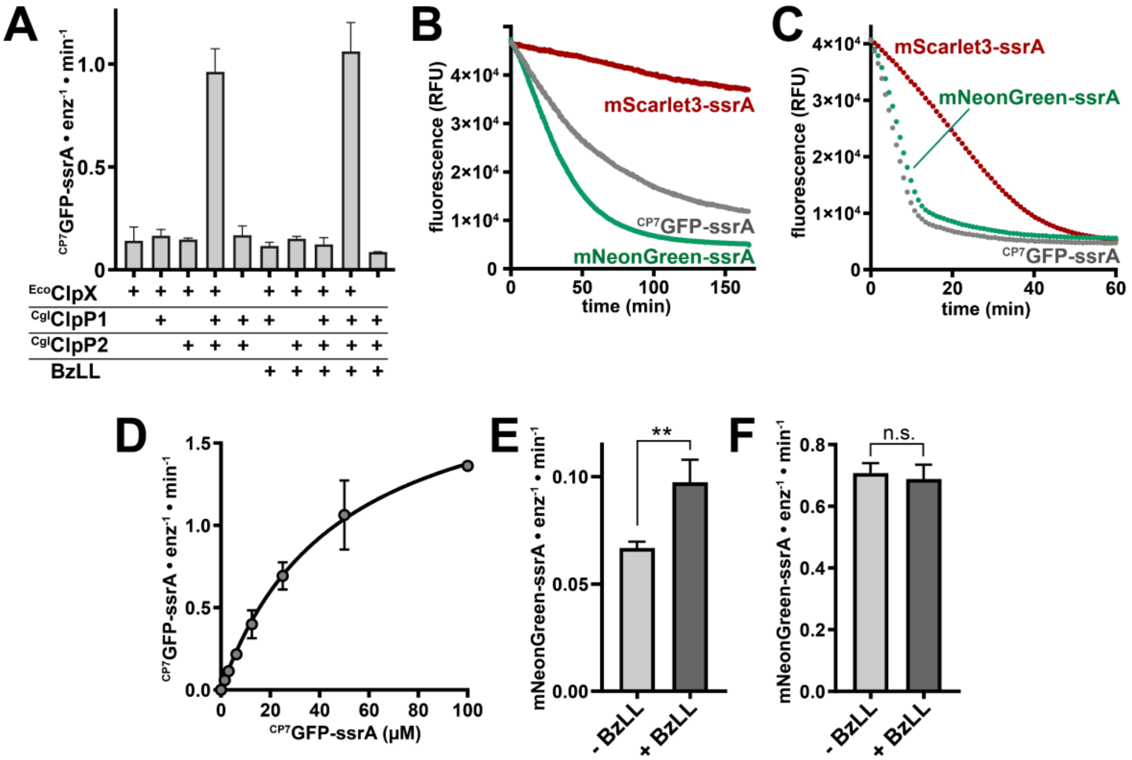

**Figure S2. Protease activity of $^{Cgl}$ClpCP1P2. A)** Proteolysis of CP7GFP-ssrA was tested with combinations of *E. coli* ClpX, $^{Cgl}$ClpP1, $^{Cgl}$ClpP2, and Bz-Leu-Leu. **B)** Trajectories follow proteolysis of the indicated substrates (20 μM) by 1 μM $^{Mtb}$ClpC1P1P2. **C)** Trajectories follow proteolysis of the indicated substrates (20 μM) by 1 μM *E. coli* ClpXP. **D)** Proteolysis of $^{CP7}$GFP-ssrA (20 μM) by $^{Cgl}$ClpCP1P2 (0.5 μM) was measured across varying substrate concentrations. Initial velocities were fit to the Michaelis-Menten equation ($K_M$ = 48 ± 7 μM, $k_{cat}$ = 2 ± 1 substrate•min$^{-1}$•enzyme$^{-1}$). **E)** Initial velocity proteolysis rates of 20 μM mNeonGreen-ssrA by 1.0 μM $^{Mtb}$ClpC1P1P2 in the presence or absence of 100 μM Bz-LL, as in Fig. 2F. **F)** Initial velocity proteolysis rates of 20 μM mNeonGreen-ssrA by 0.5 μM $^{Cgl}$ClpC1P1P2 in the presence or absence of 100 μM Bz-LL, as in Fig. 2G). Fit parameters are reported ± standard fit error. Error bars indicate standard deviation among ≥3 replicates. Significance was assessed by Student's unpaired T-test. *p*-values are indicated as follows: ** $p \leq 0.01$, n.s. non-significant.

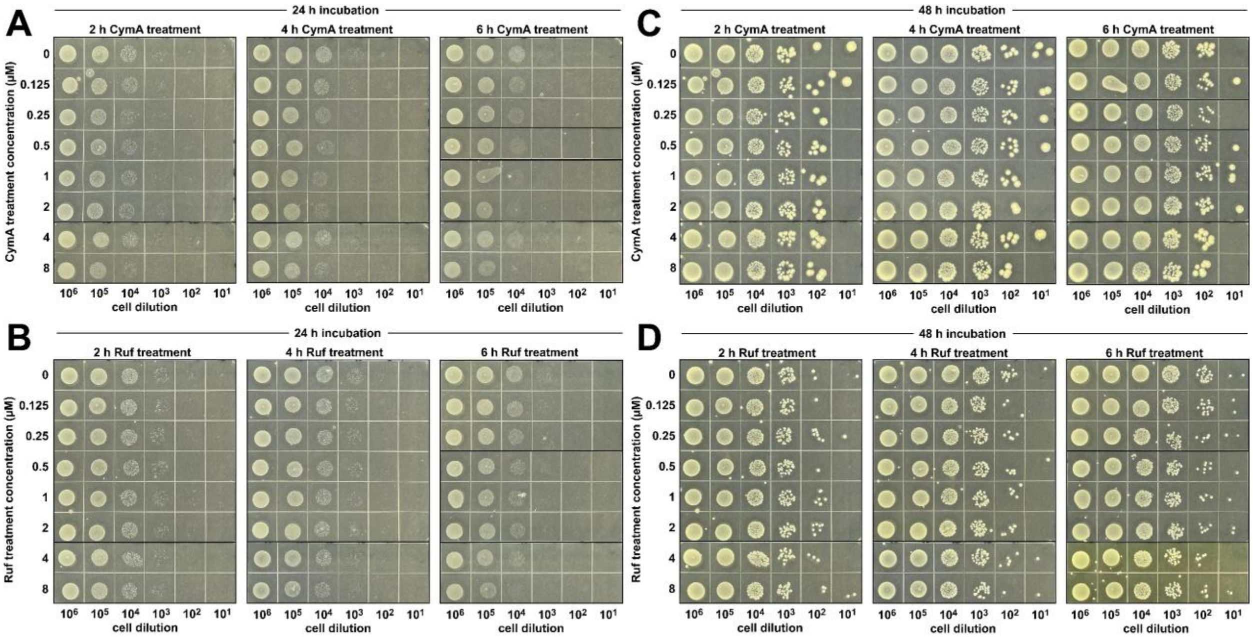

**Figure S3. Influence on CymA and Ruf on *C. glutamicum* viability.** *C. glutamicum* cultures were pre-incubated with the indicated concentrations of **A)** CymA or **B)** Ruf for varying durations, serially diluted, and spot plated on LB agar. Plates were imaged after 24 h incubation at 30°C. The same plates of **C)** CymA and **D)** Ruf treated cells were additionally imaged at 48 h.

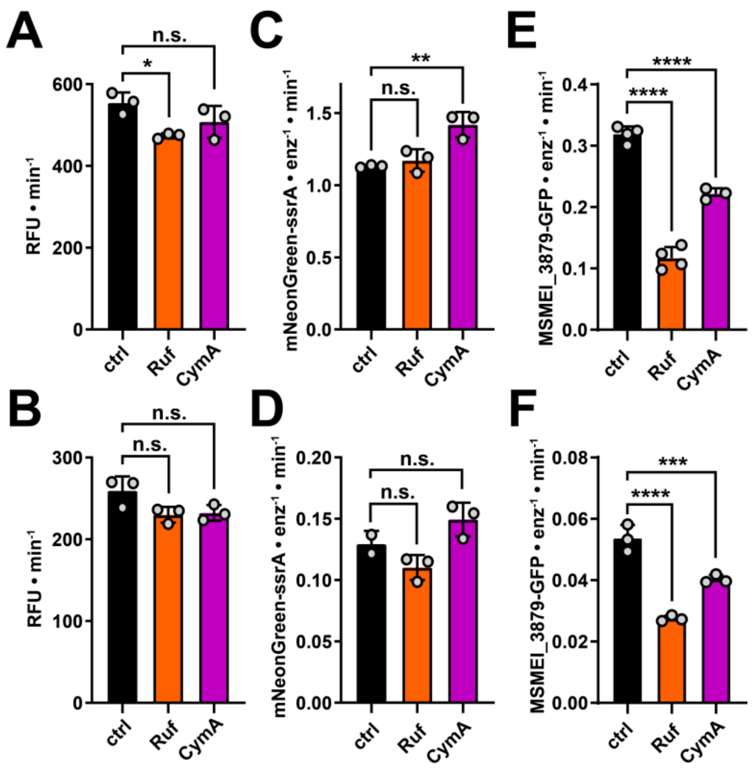

**Figure S4. Comparison of antibiotic effects on substrate proteolysis.** The maximum proteolysis rate was measured in the presence of 10 µM Ruf, 10 µM CymA, or a DMSO control on proteolysis of FITC-casein (15 µM) by **A**) $^{Cgl}$ClpCP1P2 (0.5 µM) or **B**) $^{Mtb}$ClpC1P1P2 (1 µM); mNeonGreen-ssrA (20 µM) by **C**) $^{Cgl}$ClpCP1P2 (0.5 µM) or **D**) MtbClpC1P1P2 (1 µM); or MSMEI_3879-GFP (10 µM) by **E**) $^{Cgl}$ClpCP1P2 (0.5 µM) or **F**) $^{Mtb}$ClpC1P1P2 (1 µM). Error bars indicate deviation among 3 biological replates. Antibiotic-treated samples were compared to their respective DMSO controls by ordinary one-way ANOVA followed by Dunnett's multiple comparison test. Multiplicity-adjusted p-values are indicated as follows: * p ≤ 0.05, ** p ≤ 0.01, *** p ≤ 0.001, **** p ≤ 0.0001, n.s. non-significant.
